# Castl: Robust Identification of Spatially Variable Genes in Spatial Transcriptomics via an Ensemble-based Framework

**DOI:** 10.1101/2025.10.09.681323

**Authors:** Yiyi Yu, Jiyuan Yang, Ping-an He, Xiaoqi Zheng

## Abstract

Spatially variable genes (SVGs) are essential for elucidating tissue organization within spatially resolved transcriptomics. While a number of computational methods have been developed for SVG identification, their reliance on algorithm-specific assumptions, such as predefined kernel functions or spatial neighborhood graphs, often results in substantial variability in sensitivity and inflated false discovery rates (FDRs) across heterogeneous datasets. To address this challenge, we here develop *Castl*, an ensemble-based framework for SVG identification that integrates multiple detection methods through statistically designed aggregation modules. Comprehensive evaluations on both simulated and real-world data demonstrate that Castl consistently identifies biologically meaningful spatial expression patterns, mitigates method-specific biases and effectively controls FDRs across various biological contexts, resolutions, and spatial technologies. This flexible, assumption-free framework offers a robust and standardized foundation for spatially informed feature discovery in complex biological systems.

## Introduction

The advent of spatially resolved transcriptomics (SRT) has revolutionized our understanding of gene expression landscapes within intact tissues, and provides unprecedented insights into cellular heterogeneity, tissue organization, and microenvironmental interactions (Asp et al. 2020; Armingol et al. 2021; Rao et al. 2021; Moses and Pachter 2022; Palla et al. 2022; Williams et al. 2022). One of the key analytical tasks in SRT is the identification of spatially variable genes (SVGs) (Ståhl et al. 2016), defined as genes whose expression patterns exhibit non-random spatial variation. SVGs could serve as molecular biomarkers for delineating tissue domains, inferring cell-cell communication, and uncovering spatially driven biological mechanisms underlying development, disease, and regeneration (Vickovic et al. 2019).

Recent methodological advances have led to a proliferation of SVG detection algorithms, which can be broadly categorized into kernel-based and graph-based methods (Charitakis et al. 2023; Li et al. 2023b). Kernel-based methods typically rely on spatial covariance structures defined by kernel functions, often implemented through Gaussian processes, generalized linear spatial models (GLSM), or other covariance-based statistical frameworks to model spatial dependencies. For instance, SpatialDE applies Gaussian process regression with adaptive kernel selection to capture diverse spatial expression gradients (Svensson et al. 2018); SPARK adopts GLSM to detect common spatial patterns by integrating ten different spatial kernels (Sun et al. 2020); and SPARK-X employs a non-parametric covariance-based hypothesis test to detect spatial dependence between gene expression and locations (Zhu et al. 2021). In contrast, graph-based methods generally leverage spatial neighborhood graphs constructed from coordinate information, utilizing approaches such as graph convolutional networks, autocorrelation measures, or graph signal processing techniques to capture complex spatial expression patterns. Specifically, SOMDE fits node-level representations enhanced by self-organizing maps to detect subtle spatial patterns (Hao et al. 2021); SpaGCN clusters spatial domains using graph convolutional networks and identifies domain-specific biomarkers (Hu et al. 2021); and Spanve explicitly constructs a spatial neighbor graph to quantify variation through Kullback–Leibler divergence between graph-based and random expression distributions (Cai et al. 2024).

Despite the accumulated number of SVG detection tools developed in recent years, their performance and biological interpretability are heavily influenced by dataset-specific features and underlying modeling assumptions pertaining to data distribution and spatial processes. These dependencies often lead to method-specific biases, resulting in inconsistent SVG prioritization across studies and elevated false discovery rates (FDRs) in complex tissues (Charitakis et al. 2023; Li et al. 2023b). To address these limitations, we here develop *Castl*, a computational framework that integrates multiple SVG detection methods through statistically rigorous consensus strategies. Comprehensive evaluations based on simulations and real data across a variety of resolutions, species, tissue types, and sequencing platforms demonstrate the robustness and scalability of the Castl framework in SVG identification.

## Results

### Overview of the Castl framework

Castl is a novel ensemble-based framework that integrates outputs from existing SVG detection methods to enhance SVG identification for SRT data. It currently implements three parallel aggregation strategies: rank aggregation (Rank-Agg), *p*-value aggregation (Pval-Agg), and FDP proxy-guided aggregation (FDPp-Agg) (Fig.1). Among them, Rank-Agg and Pval-Agg adopt relatively straightforward strategies, where the former computes a cumulative score derived from gene ranks and selects the top genes as SVGs, and the latter combines *p*-values of individual methods using the Cauchy combination rule (Pillai and Meng 2015; Liu et al. 2019), followed by Benjamini-Yekutieli correction (Benjamini and Yekutieli 2001) to identify spatially significant genes. In contrast, FDPp-Agg employs a more advanced approach. It first constructs an augmented expression matrix with synthetic negative controls, then computes the false discovery proportion proxy by comparing selection frequencies between controls and candidate genes. This enables adaptive determination of an optimal frequency threshold, thereby capturing subtle spatial expression signals while controlling for false positives (see Methods for details).

**Fig.1.**
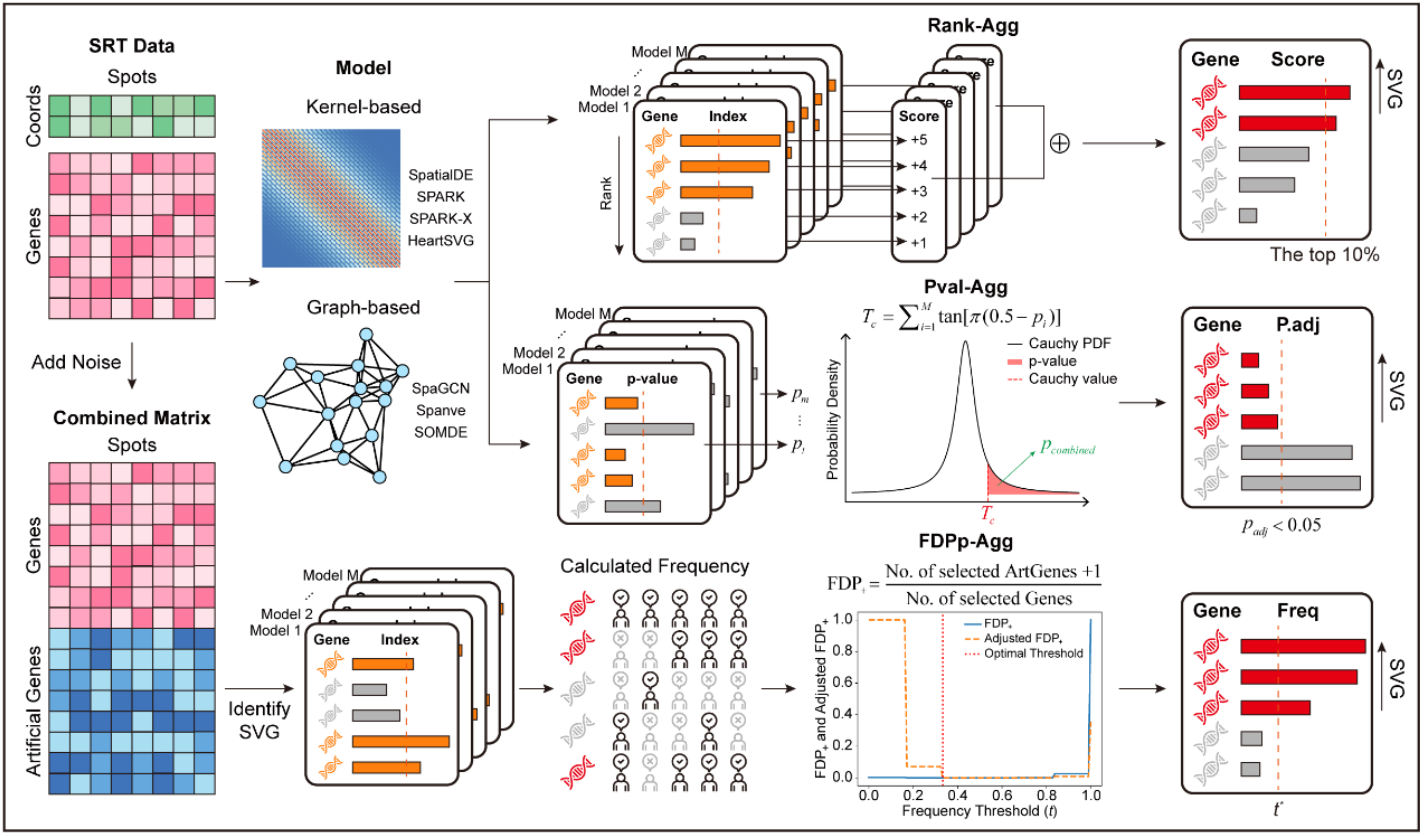
Overview of the Castl framework. Castl takes gene expression matrix and spatial coordinates as inputs. It applies multiple SVG detection methods, including kernel-based (SpatialDE, SPARK, SPARK-X, and HeartSVG) and graph-based approaches (SpaGCN, Spanve, and SOMDE), to generate baseline gene lists. It then implements three ensemble algorithms based on these gene lists: (1) **Rank-Agg**, which integrates gene rank-orders across lists into a unified composite score to identify top-ranked SVGs; (2) **Pval-Agg**, which combines *p*-values using the Cauchy combination rule followed by a multiple testing correction; (3) **FDPp-Agg**, which first generates artificial gene expressions through permutation, and then estimates selection frequencies based on combined gene expression matrix via baseline methods. The optimal threshold *t*^∗^ for SVGs is determined by minimizing the false discovery proportion with penalty adjustment.

Castl now supports seven state-of-the-art SVG detection methods as its baseline algorithms: four kernel-based methods (SpatialDE, SPARK, SPARK-X, and HeartSVG (Yuan et al. 2024)) and three graph-based methods (SpaGCN, Spanve, and SOMDE). Compared to individual methods, Castl ensures cross-method constancy while retaining the method-specific signals, thereby achieving improved identification performance and robustness in detecting complex spatial patterns. The consensus SVGs identified by Castl are used for a series of downstream data analyses, including spatial domain annotation, tissue microstructure delineation, and biomarker discovery.

### Castl enhances SVG detection performance across diverse simulated scenarios

We first evaluated the performance of the Castl framework using a series of simulated datasets generated by SRTsim (Zhu et al. 2023). To this end, we generated datasets containing both SVGs with distinct patterns (hotspots, streaks, gradients, and curves) and non-SVGs without spatial patterns using the interactive interface provided by SRTsim (Fig. 2a, Supplementary Fig. S1). Using the predetermined SVGs as a reference, we assessed the identification performance of seven baseline methods and three ensemble-based methods by calculating their F1 scores.

**Fig.2.**
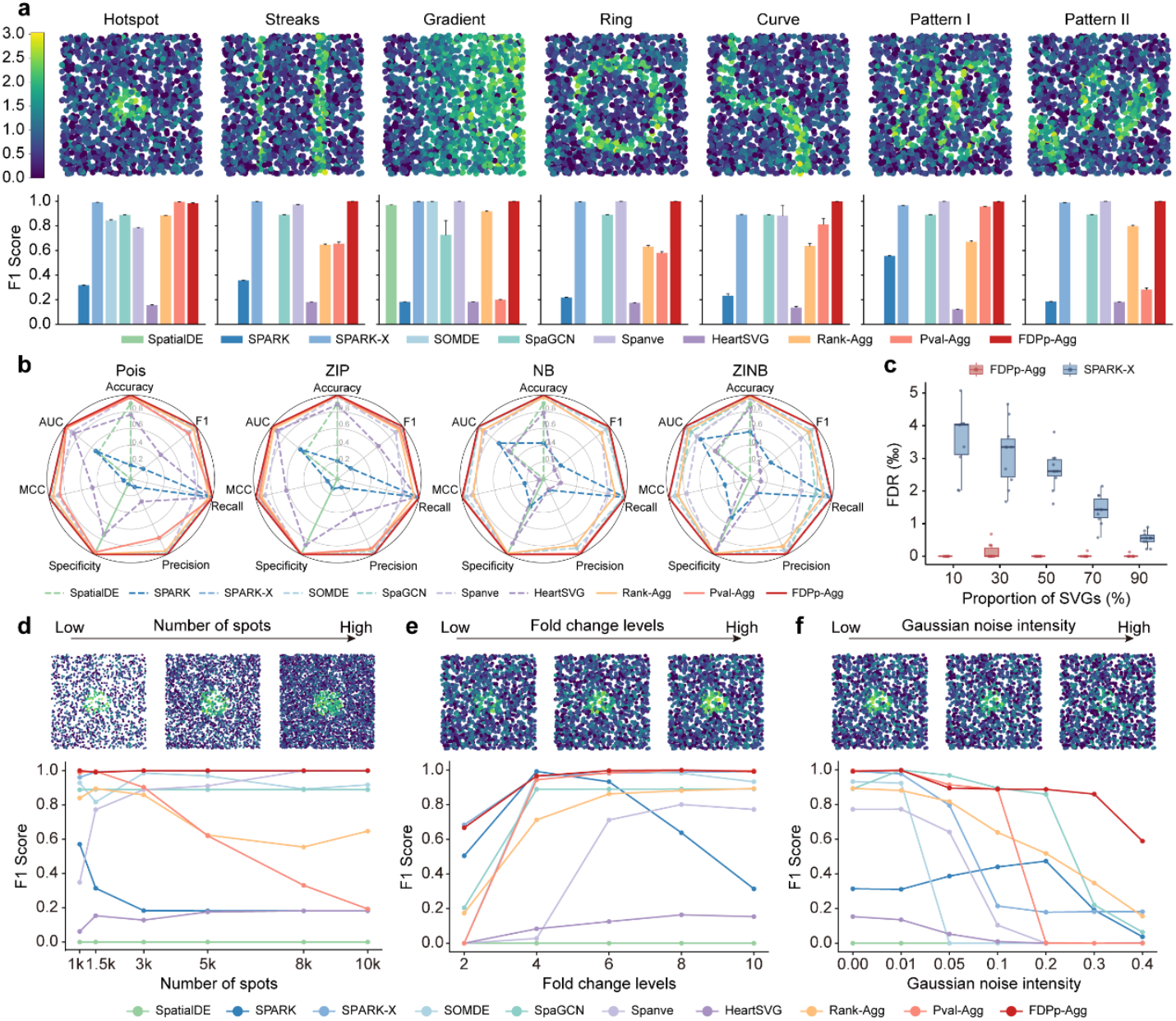
Benchmarking Castl and baseline methods across diverse simulation scenarios. **a** Visualization of simulated spatial expression patterns. Comparison of F1 scores among seven baseline SVG detection methods and three Castl ensemble-based methods. Each dataset comprises 1,500 cells and 10,000 simulated genes generated under a ZINB distribution. Error bars indicate 95% confidence intervals across 10 replicates. **b** Comparison of seven performance metrics (Accuracy, F1 score, Recall, Precision, Specificity, MCC, and AUC) for different methods in hotspot simulations under four distributions (Posi, ZIP, NB, and ZINB). **c** FDRs between FDPp-Agg and SPARK-X across varying SVG proportions. Results are obtained based on 10 replicates under ZINB distribution. **d**,**e**,**f** Visualization of gene expression patterns and F1 scores under different simulation parameters, i.e., number of spots (**d**), fold changes (**e**), and noise intensities (**f**).

As shown in Figure 2a, different baseline methods show distinct predictive performance across spatial configurations, reflecting their inherent sensitivities to specific patterns. Meanwhile, the ensemble-based methods also present distinct predictive behaviors. Rank-Agg replicates the detection trends of most baseline methods and shows a preference for broadly expressed SVGs (e.g., hotspot and gradient patterns). Pval-Agg is sensitive to both individual *p*-values and their overall distribution, but its performance becomes unstable due to numerous spuriously small *p*-values. Notably, FDRp-agg is more robust to noise and effectively compensates for biases of individual methods, achieving superior and highly stable performance (average F1 = 0.99) (Fig. 2a).

The high performance and robustness extended to different expression statistical distributions, i.e., Poisson (Pois), Zero-Inflated Poisson (ZIP), Negative Binomial (NB), and Zero-Inflated Negative Binomial (ZINB) distributions (Song et al. 2024) (Fig. 2b, Supplementary Fig. S2), and across seven evaluation metrics (e.g., F1 score, Recall, and Precision; see Methods for details). Furthermore, we examined the effect of varying proportions of SVGs among all genes. A focused comparison between SPARK-X (the best-performing baseline method) and FDPp-Agg (the most robust consensus algorithm) revealed that FDPp-Agg more effectively suppressed interference from non-SVGs, further confirming its robustness in challenging scenarios (Fig. 2c).

To further assess the scalability of the Castl framework, we varied the number of spatial spots to simulate different spot densities (Fig. 2d). Most baseline methods, including SPARK-X, SpaGCN, Spanve, and HeartSVG, show improved SVG identification accuracy with increasing spot density. In contrast, SPARK exhibits density-dependent performance degradation at high spot densities, probably due to its fixed-bandwidth kernel, which restricts adaptation to finer spatial resolution. As expected, Rank-Agg maintains consistent performance with noise resilience, whereas Pval-Agg displays instability primarily driven by SPARK’s density sensitivity. By comparison, FDPp-Agg consistently delivers robust results across all density levels (Supplementary Fig. S3), demonstrating superior adaptability to varying spatial resolutions and sampling contexts.

In addition, we evaluated Castl’s robustness under varying fold-changes (Fig. 2e, Supplementary Fig. S4) and Gaussian noise intensities (Fig. 2f, Supplementary Fig. S5). Most methods exhibit performance trends positively correlated with fold change and negatively correlated with noise intensity. Among them, FDPp-Agg achieves the highest F1 scores across all fold-change levels and noise intensities. Collectively, these comprehensive evaluations highlight the superior accuracy, robustness, and scalability superior accuracy of the Castl framework in SVG identification compared to baseline methods across diverse challenging simulated scenarios.

### Castl enables comprehensive SVG identification by leveraging complementary strengths of multiple algorithms

We next evaluated the performance of Castl on real-world SRT datasets across a variety of spatial resolutions, species, tissue types, and sequencing platforms. As a representative example, we first applied Castl to the colorectal cancer liver metastasis dataset generated by 10x Genomics Visium, which comprises four colon tissue sections and four liver tissue sections (Wu et al. 2022). Specifically, we performed unsupervised spatial clustering on the colon2 section, which contains 4,174 spots and 36,601 genes, revealing five distinct tissue domains according to the original study, i.e., smooth muscle, stromal-immune, crypt epithelium, intestinal epithelium, and tumor (Fig. 3a, Supplementary Fig. S6). The stromal-immune cluster primarily consists of stromal cells (e.g., fibroblasts) and immune cells (e.g., macrophages and T cells) within the tumor microenvironment. Of note, the ensemble-based algorithms effectively harmonize substantial variability in SVG detection across baseline methods, evidenced by the most sensitive SPARK-X (detects 13,933 SVGs) versus the most conservative SpaGCN (detects only 30 SVGs).

**Fig.3.**
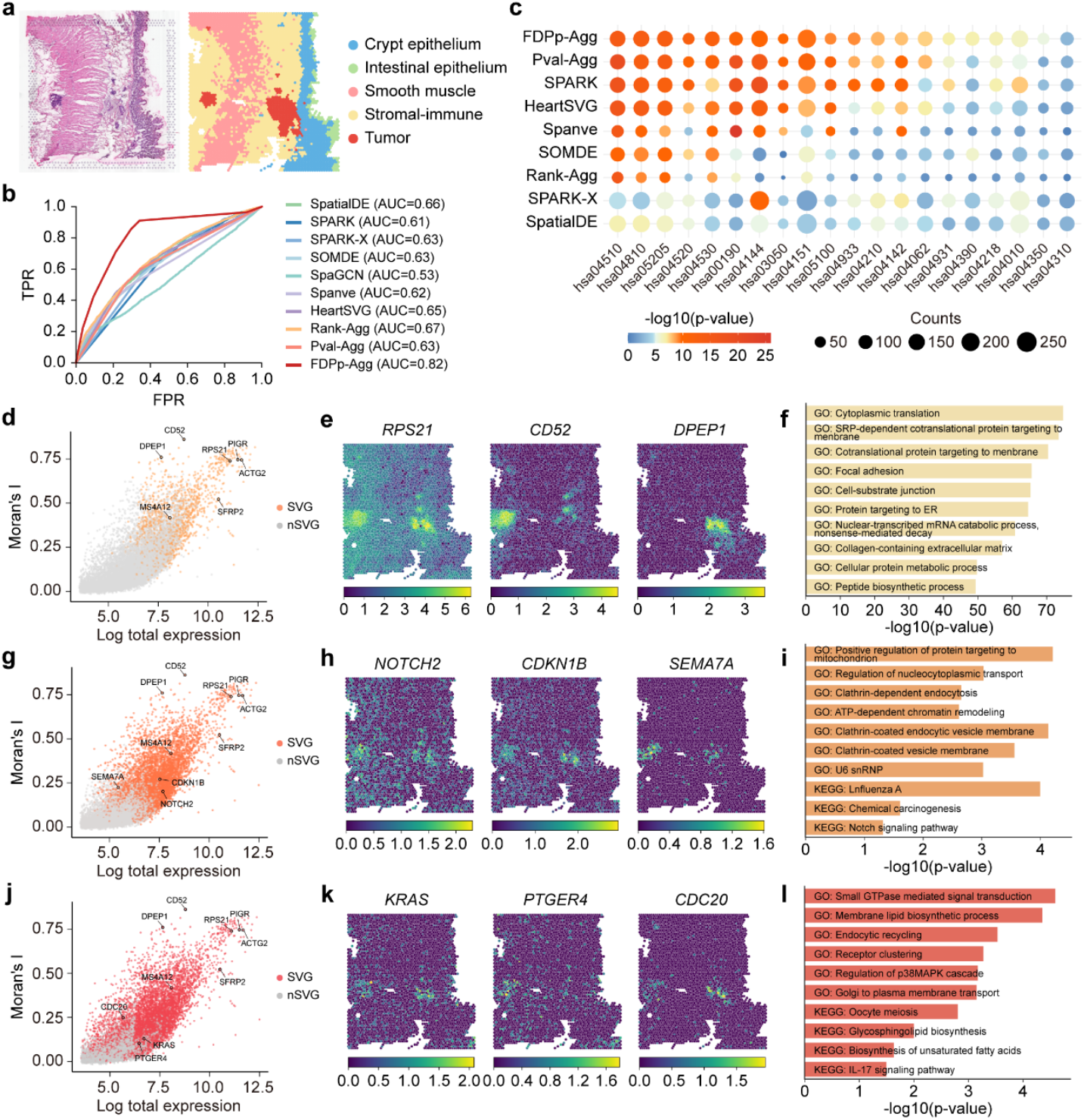
Castl improves robustness and comprehensiveness of SVGs identification in 10x Visium colorectal cancer data. **a** Hematoxylin and eosin (H&E) image of a colorectal cancer tissue section (left) and spatial clustering (right). **b** ROC curves of different methods, using gene sets from Colorectal Cancer Subtyping Consortium (CRCSC) as reference. **c** KEGG pathway enrichment of SVGs identified by different methods across key colorectal cancer pathways. Bubble size indicates overlap, color indicates significance by hypergeometric test. **d**,**g**,**j** Scatter plots illustrating spatial autocorrelation (Moran’s I) versus total gene expression across spots. Colored dots are SVGs identified by Rank-Agg (**d**), Pval-Agg (**g**), and FDPp-Agg (**j**). **e**,**h**,**k** Spatial visualizations of tumor marker genes identified by Rank-Agg (**e**), Pval-Agg (**h**), and FDPp-Agg (**k**). **f**,**i**,**l** Pathway enrichment of SVGs from Rank-Agg (**f**), Pval-Agg (**i**), FDPp-Agg (**l**), showing colorectal cancer-related pathways ranked by −log_10_ p-value.

We then benchmarked different methods by using the consensus molecular biomarkers curated by the Colorectal Cancer Subtyping Consortium (CRCSC) (Dienstmann et al. 2014; Guinney et al. 2015) as ground-truth (Roepman et al. 2014; Lee et al. 2020; Pelka et al. 2021; Schmitt and Greten 2021; Joanito et al. 2022; Qi et al. 2022). FDPp-Agg achieves the highest AUC value of 0.82, significantly outperforming both individual baseline methods and alternative ensemble approaches (Fig. 3b, Supplementary Fig. S7). Furthermore, we conducted functional enrichment analysis on SVGs identified by both individual methods and the Castl framework (Fig. 3c). Notably, SVGs identified by FDPp-Agg and Pval-Agg exhibit higher enrichment in key colorectal pathways, including cancer-related processes (hsa05200: Pathways in Cancer, has05205: Proteoglycans in Cancer), cell proliferation and apoptosis regulation (hsa04151: PI3K-Akt signaling pathway (Danielsen et al. 2015), has04010: MAPK signaling pathway (Fang and Richardson 2005), and has04210: Apoptosis), as well as cell-cell interactions and microenvironment modulation (has04510: Focal adhesion, hsa04520: Adherens junction, and hsa04530: Tight junction (Landy et al. 2016)). These findings align closely with known colorectal cancer biology, further validating the functional relevance of the identified SVGs.

To dissect the unique contributions of each ensemble-based strategy, we next analyzed their specific SVG identification patterns. Rank-Agg excels at identifying spatially autocorrelated SVGs that demarcate critical functional domains within complex tissues (Fig. 3d). Notable examples include contractility-associated *ACTG2* (Muhl et al. 2022) (Morans’I = 0.74) in smooth muscle and *SFRP2* (Andersen et al. 2024) (Morans’I = 0.52) mediating dual Wnt-immune regulation across microenvironments. Besides, it also resolves tumor heterogeneity through spatially distinct markers, e.g., a proliferative driver *RPS21* (Liu et al. 2022) enriched in tumor cores, the fibrosis-promoting *CD52* (Mennillo et al. 2024) defines stromal boundaries, and *DPEP1* (Heiser et al. 2023)-mediated inflammatory destruction shapes invasive fronts (Fig. 3e). These gene-expression patterns correspond to key morphological features, from structured stromal zones to irregular immune-infiltrated regions. Pathway enrichment analyses further confirm coordinated activation of translation, cell adhesion and extracellular matrix remodeling pathways, highlighting the efficiency of the Rank-Agg in uncovering spatial tissue architecture and progression modules of colorectal cancer.

Pval-Agg successfully identifies additional tumor-associated genes with well-established biological significance that are missed by Rank-Agg, including *NOTCH2, CDKN1B*, and *SEMA7A* (Fig. 3g,h). Notably, *NOTCH2* shows strong enrichment in Notch signaling (Wang et al. 2023), a pathway involved in tumor stem cell domain maintenance (Fig. 3i). Moreover, Pval-Agg captures fine-grained spatial structures, as evidenced by significant enrichment of subcellular localization-related pathways such as clathrin-coated endocytic vesicle membrane (GO:0030665). These findings collectively demonstrate that the statistically-driven Pval-Agg reliably identifies SVGs with critical biological functions.

Compared to two others ensemble algorithms, FDPp-Agg exhibits the broadest sensitivity spectrum, particularly in detecting low-abundance or spatially restricted genes (Fig. 3j). It successfully identifies low-expression yet highly relevant marker genes such as *KRAS* (Danielsen et al. 2015; Wood et al. 2023), *PTGER4* (Roulis et al. 2020), and *CDC20* (Ozato et al. 2023) that are frequently missed by conventional approaches (Fig. 3k). Their spatial expression patterns show significant colocalization with key pathological features, such as *KRAS* at the invasive front, *PTGER4* in immune microenvironments, and *CDC20* within proliferation hotspots. Enrichment analysis reveals that SVGs detected by FDPp-Agg are strongly associated with spatially explicit pathways (Fig. 3l), e.g., Small GTPase mediated signal transduction (Makrodouli et al. 2011) (GO: 0007264) and oocyte meiosis (has04114) reflect cellular polarity regulation and cell cycle heterogeneity at tumor margins, while membrane lipid biosynthetic process (GO: 0046467) and endocytic recycling (GO: 0032456) indicated spatial reorganization of plasma membrane microdomains. Moreover, FDPp-Agg also identifies key oncogenic genes, including *TAGLN2* (Leung et al. 2011), *GPX4* (Li et al. 2023a), and *SOX4* (Liao et al. 2008), as well as ribosomal protein genes of the RPL/RPS family (Lai and Xu 2007; Labriet et al. 2019; Liang et al. 2023) supporting the high proliferative demands of tumor cells in seven additional colon and liver sections (Supplementary Fig. S8). These results demonstrate that Castl framework achieves a more comprehensive and highly accurate identification of tumor-related spatially variable molecular markers through its complementary consensus algorithm characteristic.

### Castl enables synergistic identification of high-quality and method-specific SVGs

We next benchmarked the performance of Castl using the human dorsolateral prefrontal cortex (DLPFC) dataset generated by 10x Visium (Maynard et al. 2021). This dataset comprises 12 tissue sections from three donors with well-defined laminar architecture, including six cortical layers and white matter (Fig. 4a,b, Supplementary Fig. S9). Castl successfully identifies layer-specific marker genes such as *MOBP, TMSB10, SNAP25, PCP4, NEFM*, and *HOPX* (Fig. 4c,d), which display distinct spatial expression patterns and accurately demarcate cortical layers (Fig. 4e). In contrast, SpaGCN and Spanve are overly conservative, failing to detect *MOBP* and other functionally critical DLPFC regulatory genes with clear spatial expression patterns, such as *CARTPT, CUX2, CBLN4, NRXN1*, and *ELAVL4* (Bronicki and Jasmin 2013; Guillozet-Bongaarts et al. 2014; Hu et al. 2019; Zimmerman et al. 2020; Shibata et al. 2021; Ma et al. 2022) (Supplementary Fig. S10). Meanwhile, HeartSVG and SPARK-X identify many low-expression genes that lack spatial structure (Supplementary Fig. S11-12). In summary, Castl demonstrates strong robustness across different DLPFC sections, consistently identifying SVGs with the same spatial patterns in other section datasets (Fig. 4d, Supplementary Fig. S10).

**Fig.4.**
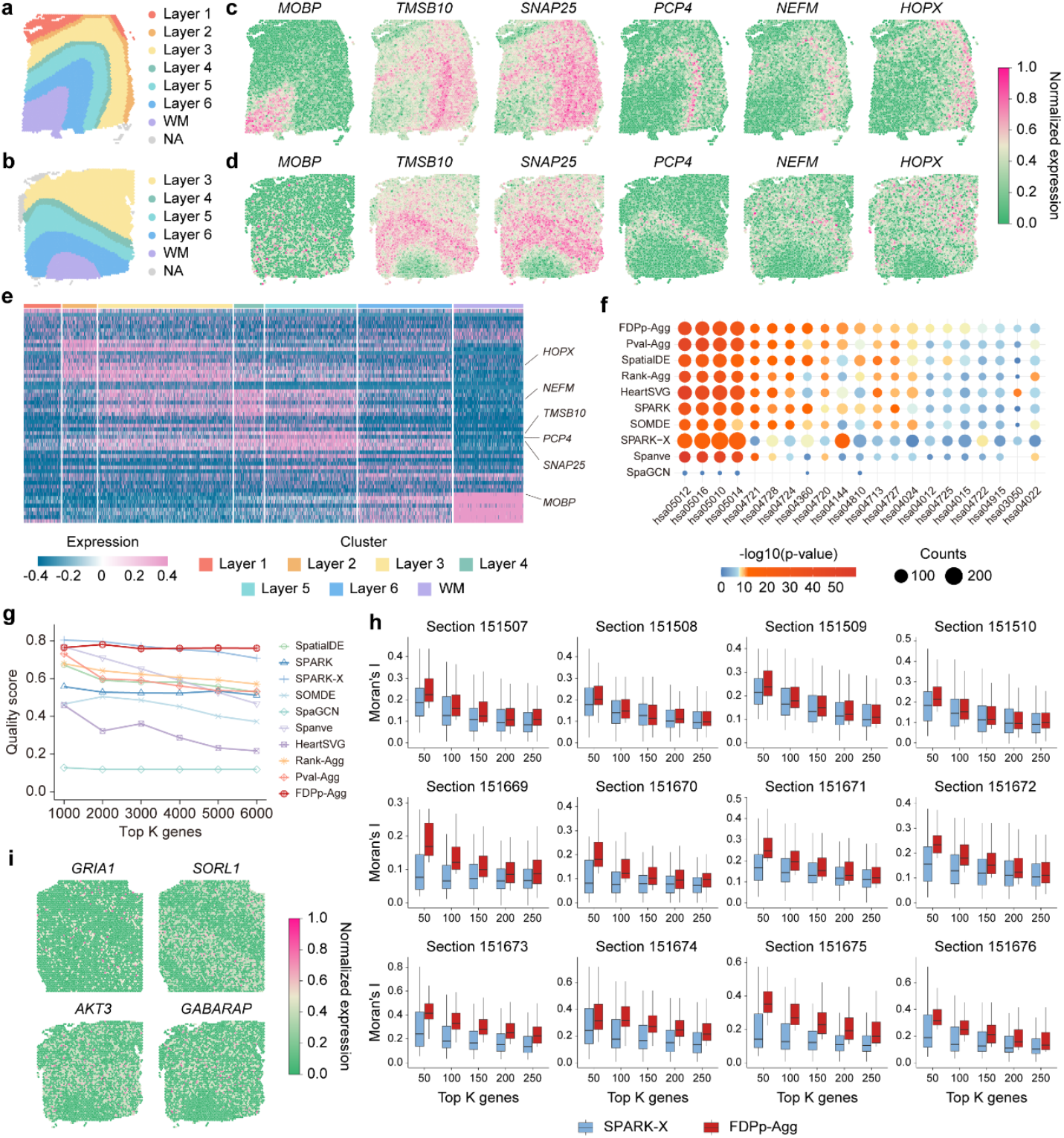
Castl enables the detection of method-specific SVGs in 10x Visium DLPFC dataset. **a**,**b** Manual annotations for sections 151673 (**a**) and 151672 (**b**) of the DLPFC dataset. **c**,**d** Visualization of representative SVGs with biological significance identified by all ensemble methods in these two sections. **e** Heatmap showing expression of layer marker genes identified by FDPp-Agg of the section 151673, representive SVGs from panel (**c**) are labeled. **f** KEGG pathway enrichment of SVGs identified by different methods across key pathways associated with cognitive function. Bubble size represents overlap, and color indicates significance. **g** Quality scores of the top SVGs by each method. **h** Moran’s I indices of the top K genes identified by FDPp-Agg and SPARK-X in each section. **i** Visualization of method-specific SVGs enhanced by FDPp-Agg.

Importantly, the SVGs identified by Castl are significantly enriched in neurodegeneration-related pathways (hsa05010 (Kumar et al. 2017) and hsa05012 (Cools et al. 2002)), synaptic function regulation pathways (hsa04724 (Moghaddam 2002) and hsa04728 (Seamans and Yang 2004)) and neuronal signaling pathways (hsa04022 (Domek-Lopacinska and Strosznajder 2005) and hsa04024 (Arnsten 2009)) (*p* < 1e-8) (Fig. 4f). Among these, FDPp-Agg identified 179 significant genes in the Alzheimer’s disease pathway (hsa05010), exceeding the average of 158 genes detected by seven other baseline methods and yielding the strongest statistical significance (*p* = 1.51e-32, Supplementary Fig. S13). These results underscore Castl’s ability to detect spatially expressed variations with high biological relevance.

To quantitatively assess the performance of different methods, we calculated the Quality Score (QS, see Methods for details), an SVG evaluation metric that integrates cross-sample consistency (technical robustness) and GO functional enrichment specificity (biological relevance, Supplementary Fig. S14). The result reveals that Castl achieves higher overall QS values than baseline methods (Fig. 4g, Supplementary Tab. S5). In particular, FDPp-Agg achieves the highest average QS (0.762), outperforming SpatialDE (0.642), SPARK (0.538), SPARK-X (0.756), SOMDE (0.460), SpaGCN (0.118), Spanve (0.643) and HeartSVG (0.313). Moreover, it exhibits remarkable stability across detection thresholds, with a coefficient of variation of just 0.3%. The top-ranked genes identified by FDPp-Agg (e.g., *GFAP, MBP, SNAP25*) display both high cross-sample reproducibility (average consistency: 0.991) and strongly enrichment in relevant GO terms, such as neuron differentiation (GO: 0030182) and synapse assembly (GO: 0007416). Spatial autocorrelation analysis using Moran’s I (Moran 1950) further confirms that SVGs detected by FDPp-Agg exhibited significantly stronger spatial clustering patterns than those identified by SPARK-X (Fig. 4h). Notably, SVGs identified by Pval-Agg achieves the highest statistically significant spatial autocorrelation (average Moran’s I = 0.316), attributed to its Cauchy combination rule (Supplementary Fig. S15). This advantage is consistently observed across all 12 DLPFC sections, validating the superior ability of Castl to detect SVGs with well-defined spatial expression patterns.

Moreover, FDPp-Agg demonstrates unique integrative capabilities by recovering method-specific SVGs that are missed by other benchmark approaches. These includes the glutamate receptor gene *GRIA1* (O’Connor and Hemby 2007) (SPARK-specific), the signaling gene *AKT3* (Danielsen et al. 2015; Chadha and Meador-Woodruff 2020; Vanderplow et al. 2021) and autophagy-related gene *GABARAP* (Piantadosi et al. 2021) (SPARK-X-specific), and the Alzheimer’s disease risk gene *SORL1* (Yu et al. 2015) (HeartSVG-specific) (Fig. 4i, Supplementary Fig. S16). Notably, none of these genes are detected by any of the six other benchmark methods, highlighting Castl’s unique ability to enhance SVG identification and recover biological meaningful genes.

### Castl resolves tissue-specific SVGs with cross-platform robustness

We next evaluated the performance of Castl on two high-resolution SRT technologies, Stereo-seq (Chen et al. 2022) and Slide-seqV2 (Stickels et al. 2021), which enable gene expression localization at subcellular to single-cell resolution. We downloaded mouse olfactory bulb (Chen et al. 2021) data generated by both platforms and annotated the resulting clusters based on anatomical references from the Allen Brain Atlas (Sunkin et al. 2013) (Fig. 5a,b). Castl successfully identifies SVGs associated with the laminar structure of the olfactory bulb across both technology platforms. The detected expression patterns show strong consistent with anatomically defined layers (Fig. 5c,d, Supplementary Fig. S17), demonstrating its robust cross-platform generalization capability. Among all methods evaluated, FDPp-Agg consistently achieves the highest enrichment significance for pathways related to neural function and olfactory regulation (Fig. 5e, Supplementary Fig. S18) and produces the highest average Quality Score for identified SVGs (QS = 0.739) (Fig. 5f, Supplementary Fig. S19 and Supplementary Tab. S6).

**Fig.5.**
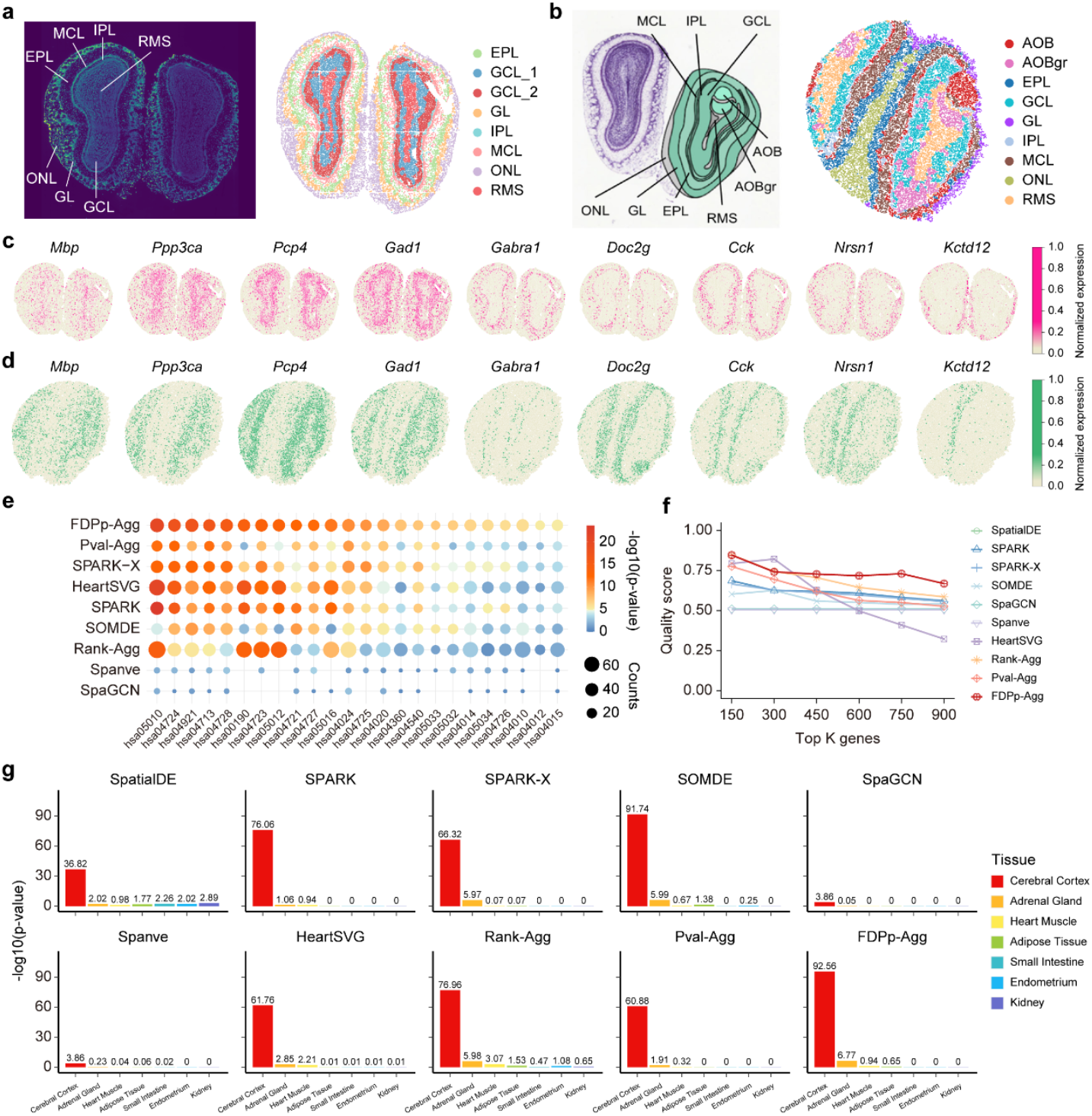
Performance of the Castl framework on single-cell resolution Stereo-seq and Slide-seqV2 mouse olfactory bulb datasets. **a** DAPI-stained image of mouse olfactory bulb laminar organization generated by Stereo-seq (left) and the corresponding spatial clustering results (right). **b** Annotations of mouse olfactory bulb laminar organization from the Allen Reference Atlas (left) and the corresponding spatial clustering results (right). **c**,**d** Visualization of SVGs associated with laminar organization as identified by all ensemble-based methods in Stereo-seq (**c**) and Slide-seqV2 (**d**) mouse olfactory bulb datasets. **e** KEGG pathway enrichment analysis of SVGs detected by different methods. **f** The top K genes and their corresponding quality scores to summarize SVG identification consistency and functional specificity of each method. **g** Tissue-specific enrichment for SVGs identified by different methods. Each subplot represents the significance of SVGs detected by each method across various tissue types.

Additionally, FDPp-Agg demonstrates exceptional tissue-specific enrichment precision (Fig. 5g, Supplementary Fig. S20). It identifies SVGs with extremely strong enrichment in cerebral cortex tissue (*p* = 2.75e-93), surpassing all other methods. This strong cortical enrichment aligns with the expected biological relevance of olfactory bulb-derived SVGs (Kawano and Margolis 1982). Furthermore, FDPp-Agg captures biologically interpretable secondary signals, showing moderate enrichment in the adrenal gland (*p* = 1.70e-7) and heart muscle (*p* = 0.115), which may reflect functional connectivity with olfactory pathways (Kobayakawa et al. 2007; Huang et al. 2015). By contrast, individual methods exhibit either limited tissue specificity or spurious off-target signals, e.g., SpatialDE shows weak enrichment in relevant tissues, Spanve and SpaGCN fail to detect meaningful signals beyond the cortex. In conclusion, these results demonstrate that Castl achieves robust tissue-specific functional associations with both high spatial precision and strong cross-platform consistency.

Finally, we extended our investigation to the single-cell resolution mouse hypothalamic preoptic region data by MERFISH (Moffitt et al. 2018). Castl robustly captures the spatial expression patterns of *Cd24a*, a key mediator of immune cell recognition and signaling (Sun et al. 2020; Yuan et al. 2024), *Prlr*, which regulates neuroendocrine hormone responses (Brown et al. 2017), and *Gabra1*, which is essential for inhibitory neurotransmission (Noriega et al. 2010). These results demonstrate that Castl can precisely identify biologically meaningful expression variation across distinct cell types (Supplementary Fig. S20-21).

## Discussion

In this study, we present Castl, an ensemble-based analytical framework that enhances the accuracy and robustness of SVG identification for SRT data by integrating outputs from multiple standard models through an ensemble-based framework. The framework is compatible with a wide range of standard models and SRT technologies, making it a versatile solution for uncovering spatially structured gene expression patterns. Comprehensive evaluations on both simulated and real-world SRT datasets demonstrate that Castl significantly improves the precision and reproducibility of SVG identification by leveraging multi-model consistency. By generating high-confidence SVG lists, Castl facilitates downstream analyses, including marker gene discovery, functional pathway enrichment, and spatial domain annotation.

Despite these advantages, Castl possesses several limitations. First, its computational complexity increases with the number of input models. Second, Castl does not explicitly account for cell-type composition or leverage cell-type deconvolution results, which may hinder accurate biological interpretation in complex tissues with heterogeneous cell populations driving spatial patterns. Third, although the ensemble approach enhances robustness, Castl remains susceptible to performance degradation if low-quality or miscalibrated models are included, underscoring the need for quality-based model weighting or filtering strategies. Finally, Castl’s reliance on a uniform spatial resolution limits its capacity to detect SVGs operating across distinct anatomical scales within heterogeneous tissues. Integrating multi-resolution datasets from the same tissue is necessary for comprehensive SVG detection, but this requires explicit handling of technical differences across spatial transcriptomic platforms. So further improvements should focus on implementing technology-aware weighting schemes to enhance cross-platform generalizability and integrating multiscale modeling or spatial hierarchy-aware approaches to accommodate varying anatomical resolutions. Incorporation of single-cell expression mapping algorithm would improve sensitivity for low-abundance transcripts. Additionally, incorporating cell-type information may improve the biological interpretation of SVGs in complex tissues and provide deeper insights into tissue-specific mechanisms.

## Methods

### Preparation of SRT data

The Castl framework takes a gene expression matrix and a spatial coordinate matrix as inputs. The gene expression matrix is denoted as *X* ∈ ℝ^*P*×*N*^, where *P* and *N* are the total number of genes and spots, respectively. Each element *x*_*ij*_ in *X* indicates the expression level of the gene *i* at spot *j*. The spatial coordinate matrix is denoted as *Z* ∈ ℝ^*N*×2^, with each element recording the spatial coordinates of the corresponding spot.

### Castl framework

Castl framework comprises three parallel ensemble modules that integrate the gene lists generated by baseline methods. Specifically,

i. Rank-Agg: Genes are ranked based on their statistical metrics (*p*-values, fold-changes, or test statistics) in a list. Each gene is then assigned an aggregation score *S*_*i*_, defined as the sum of its inverse rank across all methods. The consensus SVGs are selected as the top genes (10% by default) with the highest aggregation scores.
ii. Pval-Agg: Pval-Agg integrates multiple *p*-values (or adjusted *p*-values) from the baseline methods using the Cauchy combination rule. For each gene, the *p*-value *p*_*i,m*_ obtained from the *m*-th method are transformed into a Cauchy statistic as follows,

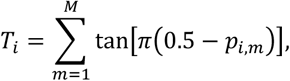

where *M* is the number of methods. The combined Cauchy statistic *T*_*i*_ is then converted into to a combined *p*-value 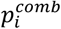. The Benjamini-Yekutieli correction procedure is applied to adjust the integrated *p*-values while controlling the FDR. Genes with *p*_*ad*j_ < 0.05 are selected as the consensus SVGs.
iii. FDPp-Agg: A **data-driven threshold optimization approach for SVG identification. It begins with a global random permutation on the original gene expression matrix** *X* to generates an artificial gene expression matrix *X*_*art*_ ∈ ℝ^*P*×*N*^ that is free of any spatial structure. The artificial gene expression matrix *X*_*art*_ is then concatenated with the original expression matrix *X* to form an augmented gene expression matrix 𝕏 = [*X X*_*art*_]^T^. The augmented combined gene matrix 𝕏 and spatial coordinate matrix *Z* are used as the baseline model inputs, yielding an extended gene list that includes artificial genes. SVGs are then calculated based on the selection frequency of each gene across all methods.

To determine a reliable frequency threshold for SVG identification, we introduced the *FDP*_4_ index:

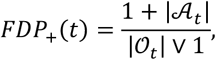

where *𝒜*_*t*_ and 𝒪_*t*_ are the number of artificial and original genes exceeding the frequency threshold *t*, and V is the maximum operator. An optimal frequency cutoff *t*^∗^ is then determined by identifying a threshold that keeps the false discovery estimate low while penalizing the inclusion of negative controls. Genes exceeding this cutoff are retained as the final consensus SVGs.

### Downstream data analysis

Based on the SVGs identified by Castl, we carried out a series of downstream data analyses to systematically assess their biological implications and the performance of the ensemble methods.

#### Differential expression analysis

Differentially expressed SVGs are selected using the *FindAllMarkers* function from the Seurat R package (Satija et al. 2015) with the thresholds |log_2_(*FC*)| > 1 and FDR < 0.05.

#### Gene enrichment analysis

Functional enrichment of significant SVGs is conducted using the clusterProfiler package (Yu et al. 2012), including the GO enrichment analysis and KEGG pathway analysis. The TissueEnrich package (Jain and Tuteja 2019) was employed to integrate multi-tissue gene expression databases, and calculate the enrichment scores for either tissue-specific SVGs or cell-type-specific SVGs.

#### Quality Score and performance evaluation

Quality Score (QS) provides a unified assessment of SVG detection methods by integrating cross-sample consistency and biological relevance. It is defined as *QS* = *C* × *FS* . Specifically, the consistency metric *C* measures cross-sample reproducibility by calculating the average detection rate of top-ranked genes across multiple biological replicates, and the functional specificity metric *FS* assesses biological meaningfulness by computing the ratio of detected genes participating in enriched GO pathways.

#### Spatial autocorrelation analysis

Spatial autocorrelation of SVGs was calculated using the Moran’s *I* index, which calculates as:

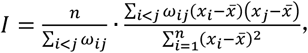

where *n* denotes the total number of spatial spots, *ω*_*ij*_ is the spatial weight indicating whether spots *i* and *j* are neighbors, and *x*_*i*_ and 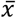 are standardized gene expression and average gene expression at spot *i* .

Statistical significance is determined through permutation tests to validate the biological authenticity of observed spatial variation patterns.

## Authors’ contributions

X.Z. and P.H. designed the study and conceived the algorithm. Y.Y. and J.Y. performed and implemented experimental analyses with the supervision of X.Z.. Y.Y., J.Y. and X.Z. contributed to the manuscript writing. All the authors have read and approved the final version of the manuscript.

## Acknowledgments

This work was supported by the National Key R&D Program of China (2024YFC2309600 to X.Z.), National Natural Science Foundation of China (62372286 to X.Z.), and Science and Technology Innovation Plan of Shanghai (23JC1403200 to X.Z). We acknowledge the Bioinformatics Core in Center for Single-Cell Omics (CSCOmics), Shanghai Jiao Tong University School of Medicine for providing bioinformatics and high-performance computing services.

## Data availability

Codes used to reproduce the results of this study are publicly available on GitHub (https://github.com/TheY11/Castl) and Zenodo (https://zenodo.org/records/16751799). All datasets analyzed in this work were obtained from the original sources. The Colorectal cancer dataset generated by 10x Visium platform is available from the Single-Cell Level Colorectal Cancer Liver Metastasis Atlas (http://www.cancerdiversity.asia/scCRLM). The human DLPFC dataset generated with 10x Visium is available from the Lieber Institute for Brain Development (LIBD) (Fan et al. 2020) (https://github.com/LieberInstitute/spatialLIBD). The Mouse olfactory bulb dataset generated by Stereo-seq and Slide-seqV2 are available from SEDR (Xu et al. 2024) (https://github.com/JinmiaoChenLab/SEDR_analyses/) and the Broad Institute Single Cell Portal (https://singlecell.broadinstitute.org/single_cell/study/SCP815/), respectively. The MERFISH dataset is available from the Dryad database (https://doi.org/10.5061/dryad.8t8s248).

